# High-dimensional proteomic analysis for pathophysiological classification of Traumatic Brain Injury

**DOI:** 10.1101/2024.04.23.590636

**Authors:** Lucia M. Li, Eleftheria Kodosaki, Amanda Heselgrave, Henrik Zetterberg, Neil Graham, Karl Zimmerman, Eyal Soreq, Thomas Parker, Elena Garbero, Federico Moro, Sandra Magnoni, Guido Bertolini, David J. Loane, David J Sharp

## Abstract

Pathophysiology and outcomes after Traumatic Brain Injury (TBI) are complex and highly heterogenous. Current classifications are uninformative about pathophysiology, which limits prognostication and treatment. Fluid-based biomarkers can identify pathways and proteins relevant to TBI pathophysiology. Proteomic approaches are well suited to exploring complex mechanisms of disease, as they enable sensitive assessment of an expansive range of proteins. We used novel high-dimensional, multiplex proteomic assays to study changes in plasma protein expression in acute moderate-severe TBI.

We analysed samples from 88 participants in the longitudinal BIO-AX-TBI cohort (n=38 TBI within 10 days of injury, n=22 non-TBI trauma, n=28 non-injured controls) on two platforms: Alamar NULISA™ CNS Diseases and OLINK^®^ Target 96 Inflammation. Participants also had data available from Simoa^®^ (neurofilament light, GFAP, total tau, UCHL1) and Millipore (S100B). The Alamar panel assesses 120 proteins, most of which have not been investigated before in TBI, as well as proteins, such as GFAP, which differentiate TBI from non-injured and non-TBI trauma controls. A subset (n=29 TBI, n=24 non-injured controls) also had subacute 3T MRI measures of lesion volume and white matter injury (fractional anisotropy, scanned 10 days to 6 weeks after injury).

Differential Expression analysis identified 16 proteins with TBI-specific significantly different plasma expression. These were neuronal markers (calbindin2, UCHL1, visinin-like protein1), astroglial markers (S100B, GFAP), tau and other neurodegenerative disease proteins (total tau, pTau231, PSEN1, amyloid beta42, 14-3-3γ), inflammatory cytokines (IL16, CCL2, ficolin2), cell signalling (SFRP1), cell metabolism (MDH1) and autophagy related (sequestome1) proteins. Acute plasma levels of UCHL1, PSEN1, total tau and pTau231 correlated with subacute lesion volume, while sequestome1 was correlated with whole white matter skeleton fractional anisotropy and CCL2 was inversely correlated with corpus callosum FA. Neuronal, astroglial, tau and neurodegenerative proteins correlated with each other, and IL16, MDH1 and sequestome1. Clustering (*k* means) by acute protein expression identified 3 TBI subgroups which had differential injury patterns, but did not differ in age or outcome. Proteins that overlapped on two platforms had excellent (*r*>0.8) correlations between values.

We identified TBI-specific changes in acute plasma levels of proteins involved in amyloid processing, inflammatory and cellular processes such as autophagy. These changes were related to patterns of injury, thus demonstrating that processes previously only studied in animal models are also relevant in human TBI pathophysiology. Our study highlights the potential of proteomic analysis to improve the classification and understanding of TBI pathophysiology, with implications for prognostication and treatment development.

## Introduction

Traumatic Brain Injury (TBI) is a highly heterogenous condition, encompassing multiple possible mechanisms and sequelae^1^. Current classifications are overly simplistic and inadequate for describing the range of processes occurring during and after TBI^2^. This limits clinical prognostication as well as patient selection for and evaluation of potential treatments. The TBI field is increasingly moving towards neuroimaging and blood biomarker-led phenotyping that can be informative about post-injury pathophysiological processes and relevant to later outcomes. Early post-TBI blood levels of neuronal and astroglial markers (e.g. NFL and GFAP) and some cytokines (e.g. IL6) reflect injury, and are associated with later neuronal loss and functional outcomes^3–5^. However, only a small number, out of likely many interacting pathophysiological processes that accompany or are triggered by TBI, have been studied.

Highly sensitive, high-dimensional protein assays are now available which can assess a very wide range of proteins using small sample volumes. These novel antibody-based proteomic technologies, namely the OLINK® Proximity Extension^6^ and the Alamar NUcleic acid Linked Immuno-Sandwich Assay (NULISA™)^7^ technologies, combine the sensitivity and specificity of immunoassay with the ability to detect a large breadth of targets^8^. This makes them ideal for discovery work, to characterise disease mechanisms and identify potential targets for intervention. This is particularly advantageous in conditions such as TBI, where a broad and complex range of processes contribute to heterogenous downstream effects.

In this study, we used the Alamar NULISA™ CNS diseases panel for the first time in a clinical TBI cohort, to investigate the plasma proteomic response in acute TBI compared with age-matched groups of non-TBI trauma (NTT) and non-injured controls (CON). The Alamar panel assesses 120 proteins, most of which have not been investigated before in human TBI. These include several phosphorylated tau species and neurodegenerative markers such as amyloid beta 42, vascular biology proteins such as VEGF-A, cytokines and proteins important for peripheral immune cell infiltration and neuroinflammation such as IL16, and proteins involved in cellular processes. This enables investigation of whether processes previously identified to be important in animal TBI models, such as autophagy^9^ and neuroinflammatory signaling^10^, are also relevant in human TBI pathophysiology. It also includes key proteins, such as GFAP and NFL, which the BIO-AX-TBI study has shown to differentiate TBI from healthy and non-TBI trauma^3^.

Samples tested were collected as part of the BIO-AX-TBI study^11^, a longitudinal TBI cohort study which also collected multiple other measures. This enabled us to investigate whether acute patterns of protein expression related to white matter injury and lesion volume in the subacute period. Further, we explored the extent to which acute patterns of plasma protein expression could help differentiate clinically meaningful TBI subgroups, since pathophysiological heterogeneity in TBI is major challenge for prognostication and developing effective intervention^1^. The explicit inclusion of an NTT group allowed us to identify specific patterns of plasma protein expression that differentiate TBI from injury in general. We additionally tested our cohorts on the OLINK Target 96 Inflammation panel to assess 92 inflammation-related proteins, and ELISA-based platforms to assess key neuronal and astroglial markers. The overlapping targets enabled us to cross-validate our findings, an important step for identifying robust protein markers for future clinical studies.

We hypothesise that (i) neuronal and astroglial markers, and proteins associated with neurodegenerative disease will be increased after TBI, whereas acute phase and inflammatory proteins will increase after both TBI and non-TBI trauma, (ii) that proteins specifically increased after TBI will correlate with injury measures from subacute MRI, and (iii) that proteomic expression can identify clinically meaningful subgroups within TBI.

## Methods

### Participant Cohort

We analysed plasma samples from n=38 TBI (33M:5F, mean age 43.8 years, *sd* 16.8), n=22 non-TBI trauma (NTT, 20M:2F, mean age 44.2 years, *sd* 17.7) and n=28 non-injured healthy (CON, 20M:8F, mean age 36.2 years, *sd* 16.2) participants. These are a subgroup of samples collected from participants in the BIO-AX-TBI study^11^, a longitudinal study of moderate-severe TBI (Mayo Criteria^12^). TBI patients mostly presented with bilaterally reactive pupils (29/35 patients), a CT Marshall grade of II (20/38 patients) (midline shift <5mm with visible basal cisterns, and no high or mixed density lesions >25cm^3^), and had a mean hospital stay of 39.3 days (*sd* 30.1) (FIG1). The Glasgow Outcome Scale-Extended at 6 months and 12 months was used to assess functional outcome after TBI^13^. The most common injury in the NTT cohort was limb fractures (FIG1E). All participants provided written informed consent, and ethical approval for the study was granted through the local ethics board.

**Figure 1:**
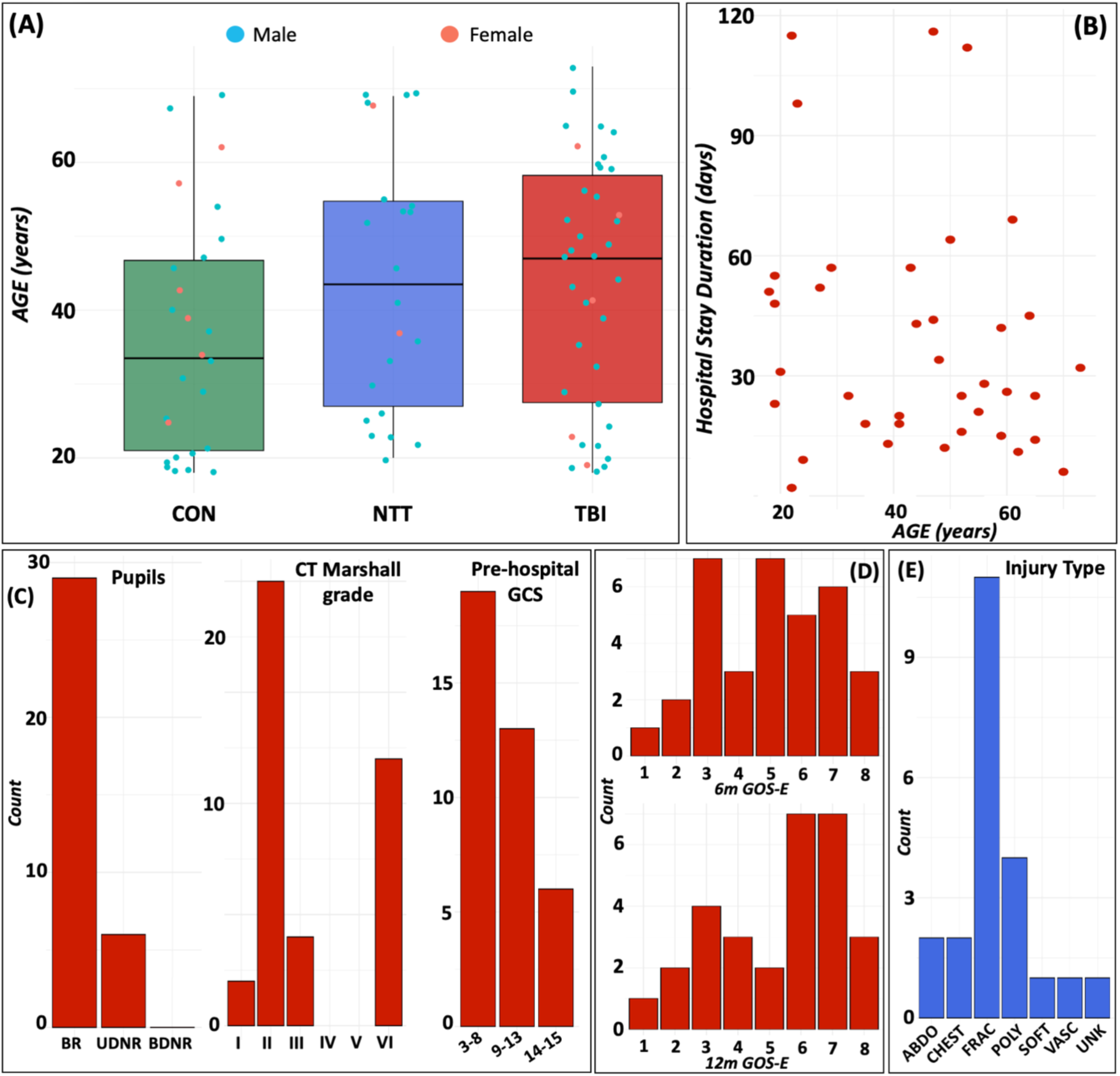
(A) Age and sex distribution of the cohorts. (B) Duration of hospital stay (days) by age for the TBI cohort. (C) Distribution of clinical descriptors of injury severity (pupil reactivity, CT Marshall grade and pre-hospital GCS) within the TBI cohort. (D) Distribution of functional outcome, assessed by Glasgow Outcome Scale Extended (GOS-E) at 6 months (6m) and 12 months (12m) after TBI. (E) Distribution of injury type within the NTT cohort.

### MRI Acquisition and Analysis

N=34 TBI (27M:7F, mean age 43.9 years, *sd* 17.2) and n=24 CON (20M:4F, mean age 35.5 years, *sd* 16.3) participants had 3T MRI scans during the subacute period (between 10 days and 6 weeks of injury) available. We included lesion volumes and measures of white matter injury in this analysis. MRI was acquired, preprocessed and analysed for the BIO-AX-TBI study (for full protocol details^3,11^). In brief, lesion volume was calculated from manually drawn lesion masks, using T1w and T2 FLAIR scans, and volumes extracted from the masks with *fslstats* from the FSL imaging analysis software package^14^. Diffusion tensor imaging was available for 29 TBI (25M:4F, mean age 43.2 years, *sd* 16.8) and 24 CON participants in the subacute period. White matter injury was assessed by calculating z-scored mean fractional anisotropy (FA) across the whole skeletonised white matter after registration of diffusion scans to DTITK space^15^ and using a tract-based spatial statistics approach to generate voxelwise maps of FA. Z-scores were calculated by comparing patients recruited on a specific scanner being compared to values from controls acquired on the same scanner. The mean z-scored FA was extracted for the corpus callosum and whole white matter skeleton. Voxelwise analysis was conducted using the general linear model with nonparametric permutation testing (10,000) in FSL Randomise^16^, with age and gender included as nuisance covariates in cross-sectional analyses and individualized lesion masking. Voxelwise analysis of zFA was cluster-corrected using TFCE, with multiple-comparison correction using a family-wise error rate of *p*<0.05.

### Blood sample processing

There were 88 plasma samples (n=38 TBI, n=22 NTT, n=28 CON) processed on the OLINK® Target 96 Inflammation panel, and 86 plasma samples (n=38 TBI, n=22 NTT, n=26 CON [20M:6F, mean age 36 years, *sd* 16.2]) processed on the Alamar NULISA™ CNS Diseases panel. Plasma samples were those which had been taken at the first acute timepoint (between day 0 and day 10 after injury in NTT and TBI cohorts) in the BIO-AX-TBI study^3^. Samples were immediately frozen at −80°C upon collection. Other plasma samples from this cohort had previously been analysed using a Simoa®-HD1 platform for GFAP, total tau, NFL and UCHL1, and using a Millipore enzyme-linked immunosorbent assay kit for S100B^3^. For this study, we tested plasma samples from the NTT cohort plus a random selection of plasma samples from the CON and TBI cohorts, after screening out those participants who had high or low outlier GFAP values on the Simoa®-HD1 platform assay. The samples were randomised onto a single plate for testing on the OLINK® Target 96 Inflammation panel^17^ and a single plate for testing on the Alamar Biotech NULISA™ CNS Diseases panel (SI Table 1). The OLINK® Target 96 Inflammation panel assesses 92 proteins implicated in inflammatory and immune pathways, whilst the Alamar NULISA™ CNS Diseases panel assesses 120 proteins associated with central nervous system diseases, including GFAP, NFL, UCHL1, S100B and total tau.

#### Alamar NULISA™ profiling

Samples used had been through 1 previous freeze-thaw cycle before analysis. Relative protein concentrations were measured by Alamar Biosciences on the NULISA™ CNS disease panel. The Alamar NULISA™ immunoassays use differential conjugation of a pair of capture and detection antibodies of each target^18^. Both antibodies are conjugated with part of a “barcode” sequence, which is specific to the target. In addition, one of the antibodies is conjugated to partially double-stranded DNA with a poly-A containing oligonucleotide, whereas the other contains a biotin-containing oligonucleotide. Sequential capture-release of the antibody/antigen and antibody/antigen/antibody complexes offers increased sensitivity, allowing a final amplification of the “barcode” sequence for each target, which is then quantified using next generation sequencing. The assay is fully automated. Normalisation takes place against both an internal control and an inter-plate control. The data for each biomarker are not absolute quantification data, but are on a log2 scale, called NULISA Protein Quantification (NPQ) units, with the data being normalised to minimise variation.

#### OLINK® protein profiling

Samples used had never been previously defrosted. OLINK Signature Q100 platform (OLINK Proteomics AB, Uppsala, Sweden) using the inflammation panel for 92 proteins. The OLINK immunoassays are based on the Proximity Extension Assay (PEA) technology^6^. Briefly, this technology uses a pair of antibodies per marker detected, labelled with oligonucleotides which are incubated with 1μl of sample. When both antibodies of a pair bind on the target, the oligonucleotides bind to each other, allowing a PCR reaction to take place, the products of which are then detected and quantified by quantitative PCR. The data for each biomarker are not absolute quantification but are given as a normalised protein expression (NPX) value. This is an arbitrary unit on a log2 scale, with the data being normalised to minimise variation.

### Statistical analysis

We carried out Differential Expression (DE) analysis on the Alamar panel data to identify proteins whose plasma levels are affects by TBI. DE analysis is a statistical approach to detect and identify, on a wide scale, biological markers whose expression varies between different groups. We used the *limma* package from *Bioconductor* in R (version 4.3.2), which uses moderated t-test, using RStudio (2021). FDR correction for multiple comparisons was made.

Spearman correlation was used to assess the correlations between blood biomarker levels identified by DE analysis, and between MRI measures of injury and those blood biomarker levels. Analyses were carried out with R (version 4.3.2) in RStudio (2021). FDR correction was used for multiple comparisons.

To investigate whether plasma protein levels could be used to identify subgroups with TBI, we performed *k means* clustering analysis using the *fmsb* package in R. We iterated from 1 to 10 k clusters, with 25 random starting assignments, and a model was built based on the k (*k*=5) which was identified as the ‘elbow’ of the scree plot of within-cluster sum of squares value, with 50 random starting assignments. The difference in age and neuroimaging measures between the clusters was interrogated using ANOVA, with cluster assignment as a factor, followed by post-hoc Tukey HSD tests if the main ANOVA was statistically significant after Bonferroni correction for multiple comparisons. Chi-squared tests were used to test whether the proportions of each GOS-E outcome category at 6 and 12 months was different between clusters.

#### Cross-validation analyses

Pearson correlations were used to determine the relationship between levels of proteins which overlapped on different assays approaches. Analyses were carried out with R (version 4.3.2) in RStudio (2021). One-way ANOVA tests (with participant type as factor) were performed for overlapping proteins using values from both available assays, in order to test whether both assay types would return the same conclusion about the main effect of cohort on protein expression.

### Data Availability Statement

the datasheets, R workspace and R code scripts will be made available to any reasonable request.

## Results

### Differential Expression analyses identify proteins associated with neurodegenerative disease, tau biology and inflammation

Differential Expression (DE) analysis of the Alamar NULISA™ CNS Diseases panel identified 71 proteins whose plasma levels were significantly different between non-injured control (CON), non-TBI injury (NTT) and TBI injury groups (FIG2A, SI Table 2). The volcano plots demonstrate how the plasma expression of proteins differed between each pair of groups (FIG2A), with the Venn diagram (FIG2B) summarising whether proteins were differentially expressed between TBI and CON only or also between TBI and NTT (FIG2B). Sixteen proteins had different plasma levels compared with both NTT and CON cohorts, indicating TBI-specific pathophysiology (FIG2A top & middle panel/2B/2C). As expected, these included neuronal (ubiquitin C-terminal hydrolase 1 [UCHL1]), astroglial (glial fibrillary acidic protein [GFAP], S100 calcium binding protein B [S100B]) and tau proteins (total levels of microtubule associated protein tau [MAPT]), which have all been previously shown to be increased after TBI compared to both CON and NTT^3^. We additionally show that changes in plasma levels of phosphorylated Tau 231 (pTau231) and amyloid beta42 (Abeta42), which have been previously seen in comparison to non-injured controls^19^, are specific to TBI and not seen in NTT (FIG2C). Newly demonstrated TBI-specific changes included deranged plasma levels of neuronal proteins (visinin-like protein 1 [VSNL1], calbindin1 [CALB1]), proteins associated with neurodegenerative disease (presenilin1 [PSEN1], 14-3-3γ [YWHAG]), inflammatory signalling proteins (ficolin 2 [FCN2]) and proteins involved in cell signalling (secreted frizzle-like protein1 [SFRP1]), cell metabolism (malate dehydrogenase 1 [MDH1) and autophagy (sequestosome-1 [SQSTM1]). Whilst most of these proteins had significantly raised levels in acute TBI, the levels of FCN2, Abeta42 and SFRP1 were significantly lower in TBI than the CON and NTT cohorts (FIG2B/C).

**Figure 2:**
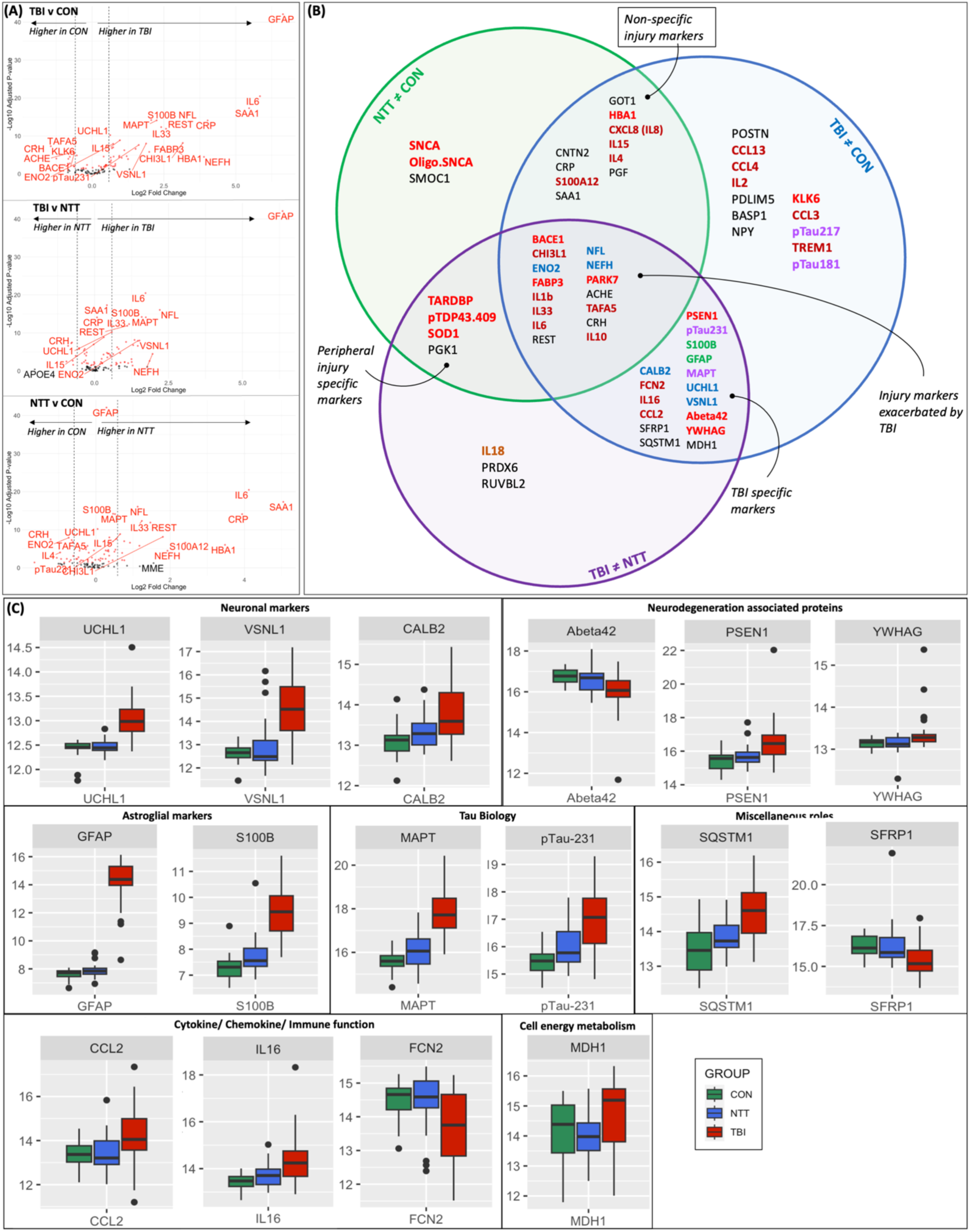
(A) Volcano plots showing the differences in plasma protein expression between each group pair. Red dots denote significant proteins. (B) Schematic of the categorization of proteins with significant group differences (adjusted p<0.05). Proteins are categorized based on a between-group comparison coefficient log_2_>0.58 (equivalent to >1.5x difference between the two groups) and post-hoc t-test p<0.05. Protein names colour-coded based on biological role/pathway: red=neurodegenerative disease associated; blue=neuronal marker; orange=cytokine/chemokine; purple=tau pathology; green=astroglial marker. Black indicates a range of other roles and pathways (SI TABLE). Nine proteins where between-group comparison did not meet these criteria for any of the group pairs (TBI v CON, TBI v NTT and NTT v CON) are not included in the schematic. (C) Boxplots illustrating plasma protein levels in CON, NTT and TBI groups for proteins identified by DE analysis to TBI specific. Y-axis units are NPQ.

A number of proteins were raised in both NTT and TBI groups, but more so after TBI, suggesting they have a role in the pathophysiology of general injury that is exacerbated by TBI (FIG2B/C, SI FIG1). These included pro-inflammatory cytokines (interleukins 1b, 33, 6), anti-inflammatory cytokines and proteins (interleukin 10 and chitinase-3-like 1 [CHI3L1]), as well as neuronal proteins (neurofilament light [NEFL] and enolase 2 [ENO2]). Non-specific injury markers, that is proteins with increased plasma levels in the NTT group with no additional effect of TBI, included acute phase proteins (C-reactive protein [CRP], serum amyloid A1 [SAA1]), and inflammatory proteins (e.g. interleukin 8) (FIG2A bottom panel, SI FIG1).

### Correlations between acute levels of TBI-specific proteins and subacute MRI findings

Diffusion tensor imaging was used to quantify white matter injury. Compared to the CON cohort, the TBI cohort had significantly reduced white matter fractional anisotropy (FA) z-scores across the both the whole skeletonised white matter tract and the corpus callosum in the subacute setting (scanned 10 days to 6 weeks after TBI), indicative of white matter injury (FIG3A). Tract Based Spatial Statistics analysis also showed significant reductions in FA present in several white matter tracts (FIG3B). Further, there was an inverse relationship between acute plasma levels of the cytokine CCL2 and corpus callosum FA *z*-scores (r_s_=-0.60, *p*=0.0006), and a positive relationship between acute levels of the autophagy related protein sequestome1 (SQSTM1) and (r_s_=0.5, *p*=0.0054).

**Figure 3:**
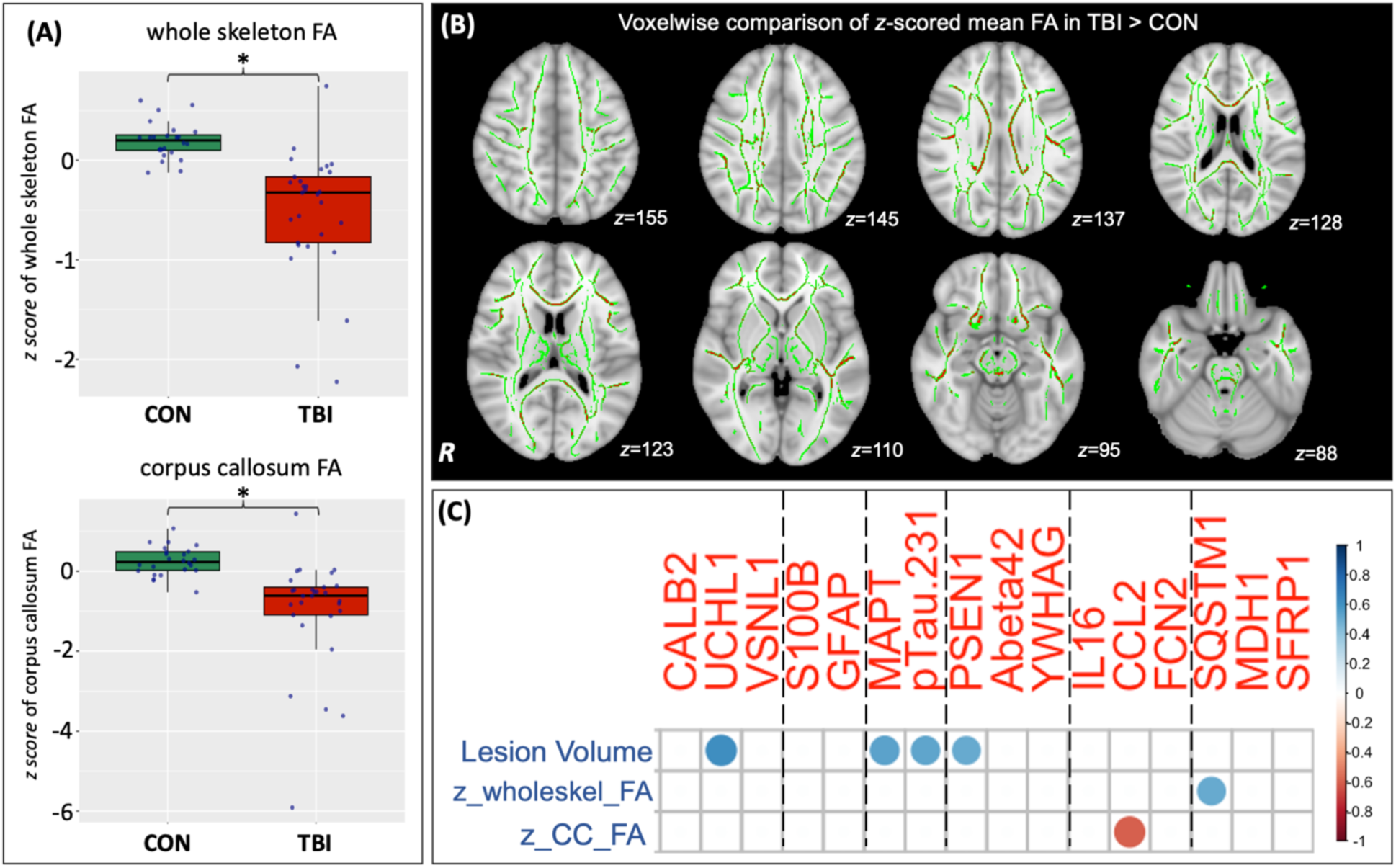
(A) Comparison of mean Fractional Anisotropy (FA) z-scores of the whole skeleton and corpus callosum between non-injured healthy controls (CON) and TBI patients on subacute MRI (10 days to 6 weeks post-injury). *denotes adjusted p<0.05. (B) Voxelwise comparison of z-scored mean FA in patients compared to controls on subacute MRI (10 days to 6 weeks post-injury), with significant (p<0.05) group differences in red, overlaid on the white matter skeleton in green. Results are overlaid on a 1mm standard brain in DTITK space. (C) Correlation matrix of TBI-specific proteins (identified by DE analysis) with lesion volume (Lesion Vol), z-score of the whole skeleton FA (z_wholeskel_FA) and z-score of the corpus callosum (z_CC_FA) on subacute MRI (10 days to 6 weeks post-injury) within the TBI cohort. Proteins ordered by category, separated by dashed line. See SI Table 2 for category. Only statistically significant results (FDR adjusted p<0.05) are shown.

Thirty-two TBI patients had evidence of focal lesions (mean volume=22,198.55mm^3^, range=352.00–82147.05mm^3^). Subacute lesion volume was positively correlated with acute plasma levels of the neuronal marker UCHL1 (r_s_=0.64, *p*<0.0001), the amyloid cleavage enzyme presenilin 1 (PSEN1) (r_s_=0.53, *p*=0.0012), total tau (MAPT) (r_s_=0.61, *p*<0.0001) and the phosphorylated tau isoform pTau231 (r_s_=0.62, *p*<0.0001), indicating a potential relationship to extent of focal injury (FIG3C).

### Correlations between acute levels of TBI-specific proteins

There were multiple significant correlations between the TBI-specific proteins, particularly between neuronal and astroglial markers, tau proteins and proteins associated with neurodegeneration (FIG4). In turn, many of these proteins showed significant correlations with acute plasma levels of IL16, sequestome 1 (SQSTM1) and malate dehydrogenase 1 (MDH1). Whilst most were positive correlations, MDH1 levels were inversely correlated with levels of plasma Abeta42, while FCB2 was inversely correlated with pTau231 but positively correlated with Abeta42.

**Figure 4:**
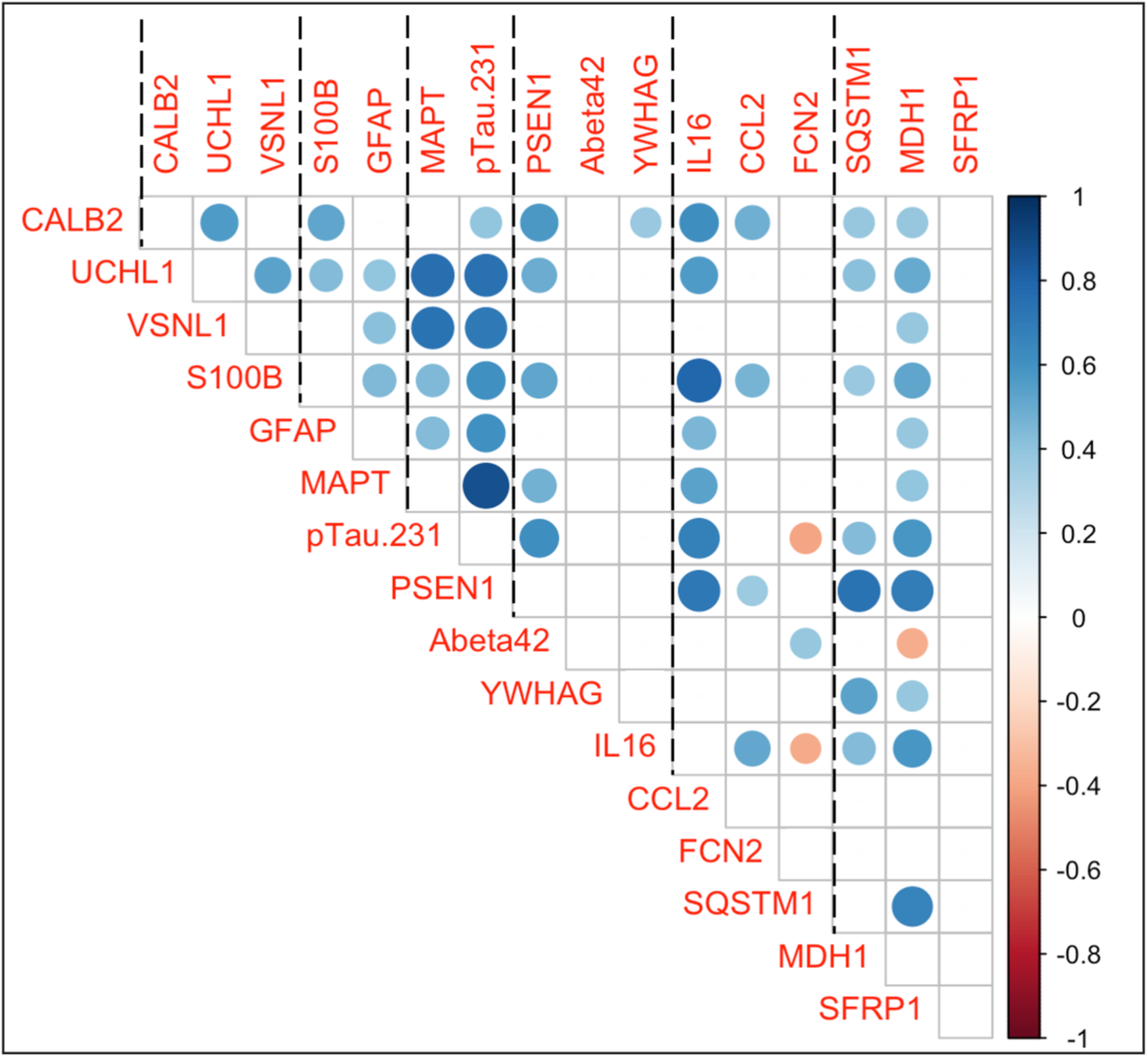
Correlation matrix of TBI-specific proteins (identified by DE analysis) within the TBI cohort. Proteins ordered by category, separated by dashed line. See SI Table 2 for category. Only statistically significant results (FDR adjusted p<0.05) are shown.

### Cluster analysis of acute plasma proteins identifies distinct TBI subgroups with differing subacute MRI findings

We next performed clustering analysis to assess whether variation in plasma protein levels could identify distinct groups in a data-driven way. *K* means cluster analysis of plasma proteins using the Alamar NULISA™ CNS Diseases panel identified 5 groups (total wiss=6988.2, between SS/total SS=31.5) (FIG 5A). We were able to separate TBI groups (cluster 3, 4, 5) from non-TBI groups (clusters 1, 2). Further, this analysis separated the TBI cohort into subgroups, in particular, differentiating a group with a high burden of focal injury (cluster 4) (FIG 5A/B right panel). There was a main effect of cluster type on mean skeleton FA *z-*score (F(4)=6.624, p=0.00025), corpus callosum FA *z-*score (F(4)=7.151, p=0.000134) and lesion volume (F(4)=10.33, p=2.45e^-06^) assessed on subacute MRI (FIG5B). Post-hoc Tukey tests confirmed that TBI cluster 5 had lower corpus callosum and mean skeleton FA *z-*scores than both non-TBI clusters (clusters 1, 2), whilst TBI cluster 4 had greater lesion volumes than both non-TBI clusters and TBI cluster 5 (FIG5B).

**Figure 5:**
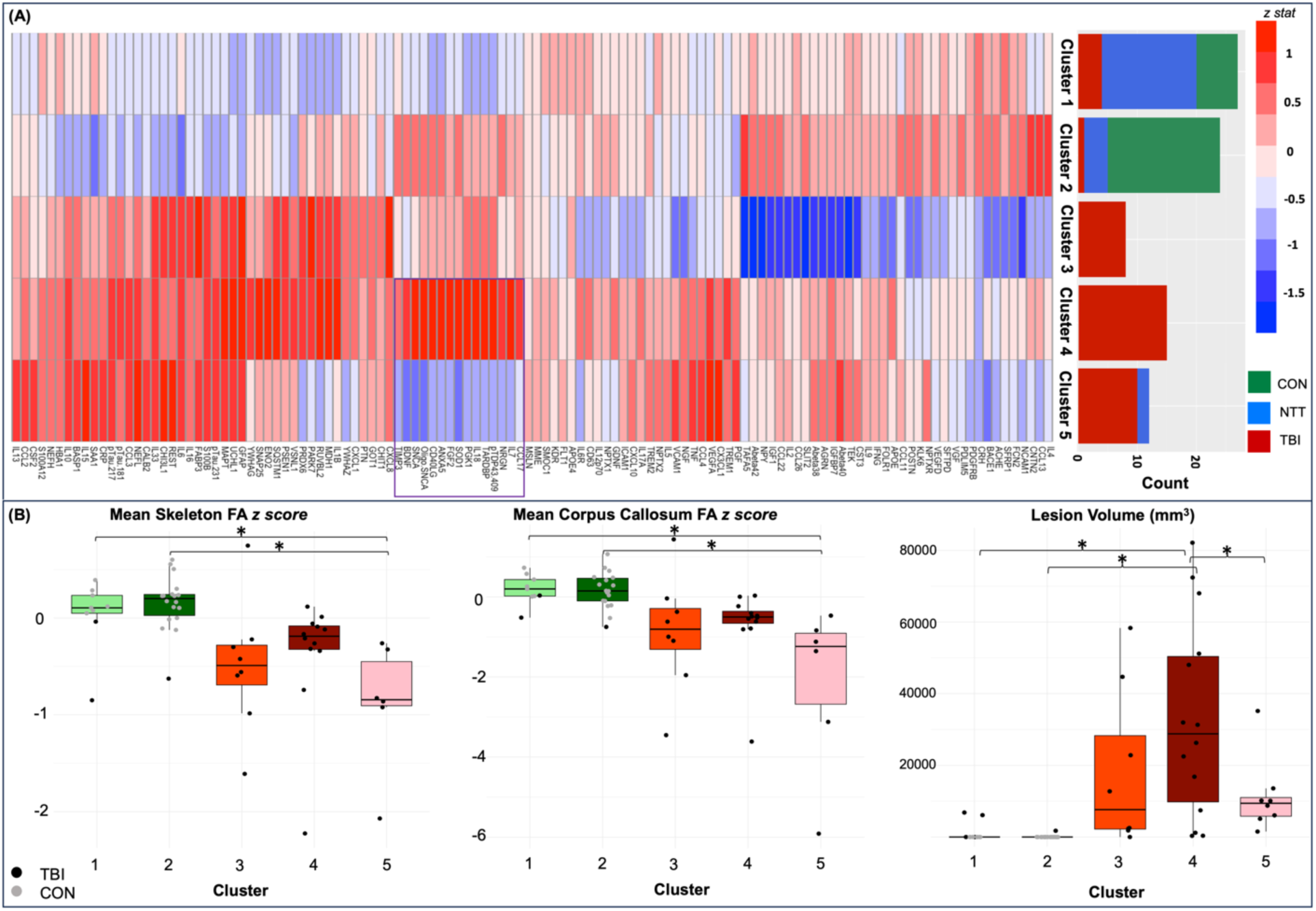
(A) Cluster analysis identified 5 clusters, with 3 TBI-predominant clusters (3,4,5) and 2 non-TBI predominant clusters (1,2). Purple box highlights a set of proteins with marked differently levels between TBI-predominant Clusters 4 and 5. CON=non-injured healthy controls, NTT=non-TBI trauma controls, TBI=traumatic brain injury. (B) Comparison of MRI measures between the clusters. Cluster 5 had significantly lower mean skeleton FA and mean corpus callosum FA, compared to Clusters 1 and 2. On the other hand, Cluster 4 had significantly higher lesion volumes than Clusters 1,2 and 5. *denotes p<0.05 on post-hoc Tukey test, performed after a statistically significant effect of cluster was identified using ANOVA. FA=fractional anisotropy. Note that only CON and TBI groups had MRI.

**Figure 6:**
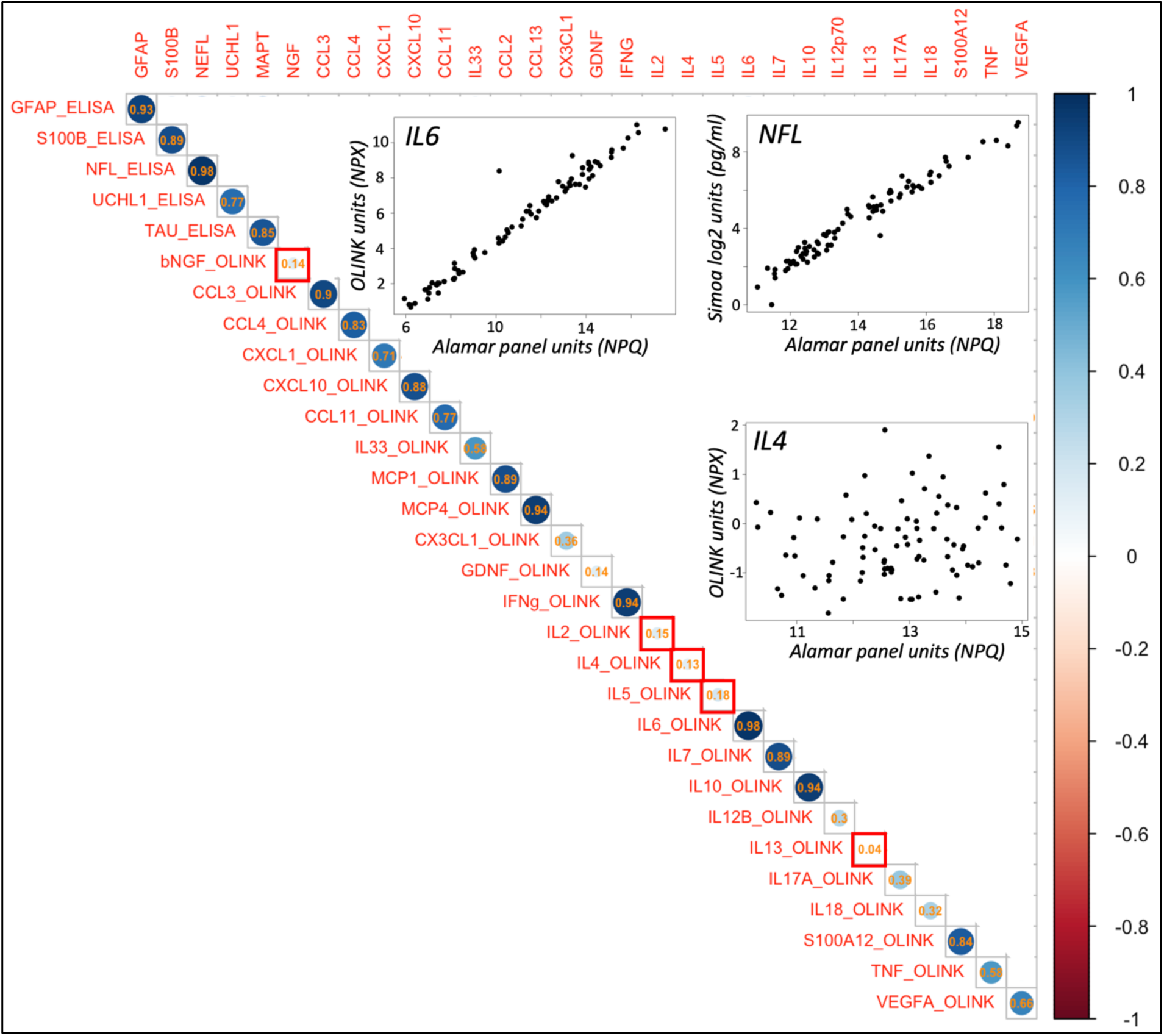
Pearson correlation between levels of proteins assessed on two different assay approaches (the Alamar NULISA™ CNS Diseases panel, horizontal, versus OLINK® Target 96 Inflammation panel or Simoa® or Millipore ELISA-based assay, vertical). Correlation coefficient is shown superimposed on representative circle. Red boxes denote proteins were >50% samples were below the limit of detection (LOD) of the OLINK panel. We used 50% of samples as the cut-off because this would mean that below the LOD would not simply be from one participant type. Inset are plots of three of the overlapping proteins: IL6 and GFAP, which show excellent correlation between two assay approaches, and IL4 which shows very poor correlation. (Note: NEFL/NFL, MAPT/total TAU, MCP1/CCL2 and MCP4/CCL13 are the same proteins, but named differently on the different assays).

**Figure 7:**
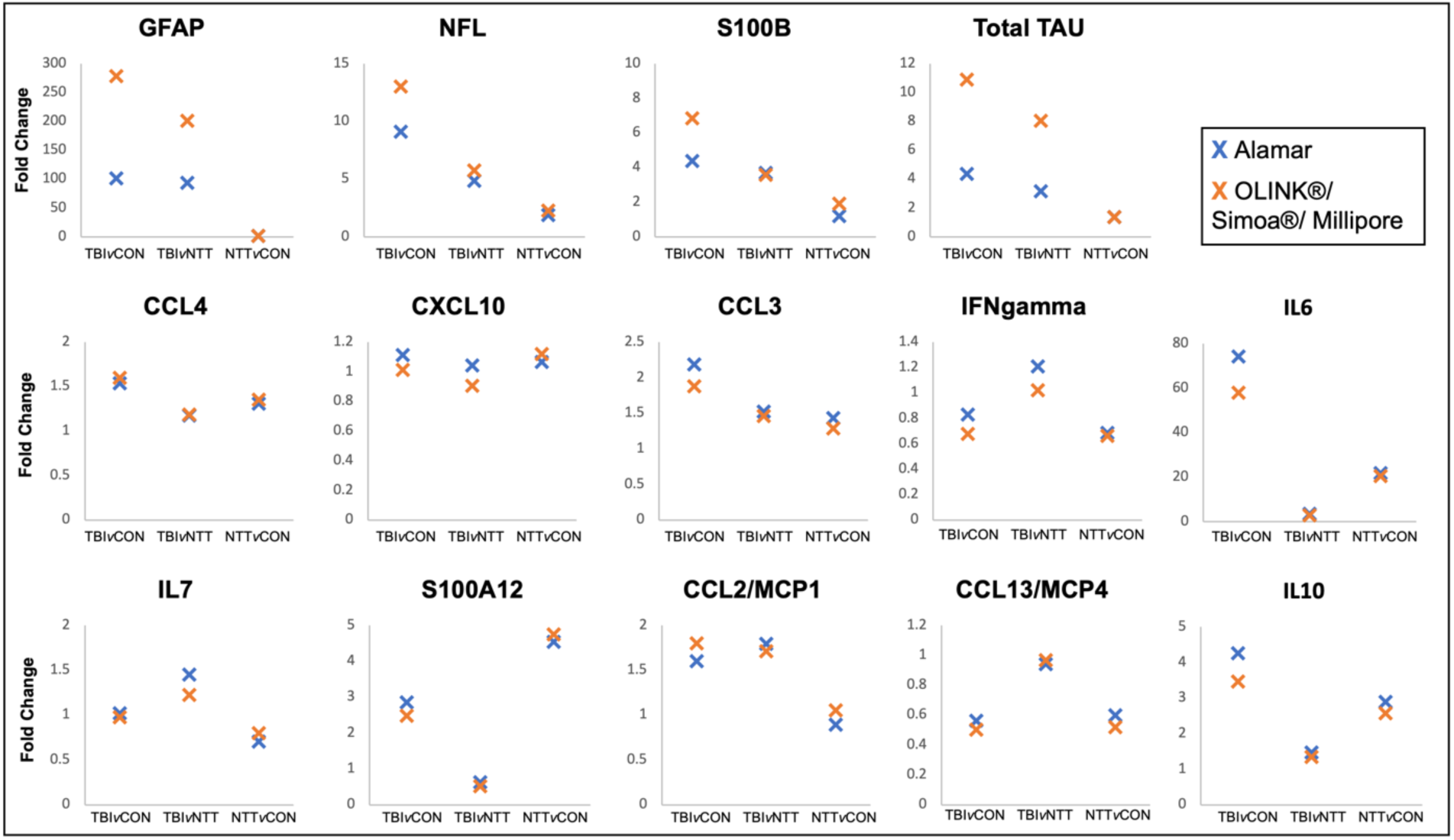
Fold differences for proteins assessed on two assays, where the correlation between levels of proteins assessed on the assays >0.8. TBIvCON denotes comparison between TBI and CON cohorts, TBIvNTT denotes comparison between TBI and NTT cohorts, NTTvCON denotes comparison between NTT and CON cohorts.

Compared to control-predominant clusters (Cluster 1 and 2), the TBI-predominant Cluster 5 had significantly lower mean skeleton FA z score (mean difference between Cluster 5 and Cluster 1= - 0.92, 95% CI[-1.67, −0.17], p=0.009; mean difference between Cluster 5 and Cluster 2=-1.03, 95% CI[-1.70, 0.37], p=0.0006) and corpus callosum FA z score (mean difference between Cluster 5 and Cluster 1=-2.33, 95% CI[-3.86, −0.80], p=0.0007; mean difference between Cluster 5 and Cluster 2= −2.30, 95% CI[-3.67, −0.93], p=0.0002). On the other hand, the TBI-only Cluster 4 had significantly greater lesion volumes than the control-predominant Cluster 1 (mean difference=31672.74, 95% CI[13559.48, 49786.00], p=0.00007) and Cluster 2 (mean difference= 32761.78, 95% CI[17096.16,48427.40],p=0.000002). Furthermore, Cluster 4 also had significantly higher lesion volume than Cluster 5, despite both being TBI groups (mean difference=21545.40, 95% CI[1620.72,41490.90],p=0.029). Plasma proteins whose acute levels were higher than the whole group mean in cluster 4, and lower than the whole group mean in cluster 5, included inflammasome-associated proteins (e.g. interleukin 18), proinflammatory cytokines (e.g. interleukin 7) and proteins associated with neurodegenerative disease (e.g. alpha-synuclein [SCNA], oligo-alpha-synuclein [SCNA], superoxide dismutase [SOD1], TAR DNA binding protein 43/TDP-43 [TARDBP] and TDP-43 with phosphorylation on serine 409 [pTDP43.409]) (FIG5B). There was no difference in age (F(4)=1.251, p=0.296) or in GOS-E at 6 months (X-squared = 2.1325, df = 4, p-value = 0.7114) or 12 months (X-squared = 2.902, df = 4, p-value = 0.5744) between any of the clusters.

### Proteins measured with two assays showed excellent overall agreement

As a range of proteomic platforms become available, an important issue is whether results are consistent across panels. We performed Pearson correlations for proteins which were assessed on two different assays (Alamar NULISA™plus either OLINK®/ Simoa®/ Millipore), and found high correlation coefficients (r>0.8) for the majority of overlapping proteins (FIG6). Notably and unsurprisingly, the proteins where >50% of samples were flagged as below the limit of detection on one of the panels (neuronal growth factor beta-subunit, IL2, IL4, IL5 and IL13), had very low correlation coefficients. Additionally, for all proteins which were tested on 2 assays with a correlation coefficient >0.8, ANOVA testing drew the same conclusion about effect of group (CON, NTT and TBI) on protein levels (SI FIG 1). Additionally, all but two proteins (MCP1/CCL2 and IFNγ) had the same group differences on post-hoc testing (SI FIG1, SI Table 3).

### Fold changes detected show differences between assay approaches

Proteins assessed on both the Alamar NULISA™ and ELISA-based platforms (Simoa® or Millipore), and where the correlation between assay values was >0.8, were: GFAP, S100B, NFL and total TAU. For these proteins, ELISA-based approaches detected larger fold changes compared to the Alamar NULISA™ panel particularly between TBI and CON cohorts (FIG7). Proteins assessed on both Alamar NULISA™ and OLINK® assays, with correlation between values >0.8, were: CCL4, CXCL10, CCL3, IFNgamma, IL6, IL7, S100A12, CCL2/MCP1, CCL13/MCP4 and IL10. The platforms detected similar fold changes in these proteins. The OLINK® Target 96 Inflammation assay detected slightly larger fold changes in CCL2, CCL4 and whilst the Alamar NULISA™ CNS Diseases assay detected slightly larger fold changes in the other proteins, in the TBI versus CON comparison.

## Discussion

Using the novel multiplex proteomic assay, Alamar NULISA™ CNS Diseases, we identified several proteins whose plasma expression were altered specifically in acute TBI patients, compared to non-TBI trauma and healthy participants. We show TBI-specific deranged plasma levels of proteins associated with neurodegenerative disease (PSEN1, 14-3-3γ), immune signalling (FCN2, SFRP1), cell metabolism (MDH1) and autophagy (SQSTM1). Thus, we provide in human evidence of several pathways being important for TBI pathophysiology, which have previously only been shown in animal studies. Additionally, we show that changes in plasma phosphorylated Tau-231 (pTau231) and amyloid beta-42 are specific to TBI, and not NTT. We further replicate, on a novel multiplex immunoassay platform, previous findings that neuronal and astroglial markers (GFAP, S100B, UCHL1, TAU and NFL) have utility as TBI-specific markers. Conversely, our study suggests that increased plasma levels of some proteins, such as IL6 and IL10, may be a non-specific response to any injury. We related acute changes in TBI-specific plasma proteins with each other, and acute changes in plasma levels of UCHL1, total TAU, pTau231, PSEN1, CCL2 and SQSTM1 with subacute neuroimaging measures of injury. Further, we found that different patterns of plasma protein expression can identify TBI subgroups that have specific injury pathologies. Overall, we show that studying acute patterns of plasma protein expression can help quantify focal injury and identify potential processes, including neurodegeneration and inflammation, that are important in human TBI pathophysiology. This provides a basis for more rational classification of TBI based on pathophysiology, We found acutely raised levels of inflammatory proteins IL6, IL10, IL1b, IL8, IL15, IL16, serum amyloid A (SAA1) and CCL2 after TBI, in line with prior studies^20,21,30–34,22–29^. In contrast to many prior studies, we recruited a non-TBI trauma control cohort, and thus demonstrate that changes in plasma levels of IL8, SAA1 and IL15 may actually reflect a general injury response. IL1b, IL10 and IL6 levels were raised in both NTT and TBI compared to CON, but were higher in TBI than NTT, suggesting that these cytokines reflect a general proinflammatory response to injury that is exaggerated by TBI.

Conversely, IL16 and CCL2 were specifically elevated in TBI, which may reflect their roles in promoting neuroinflammation. CCL2 is a key mediator of post-injury neuroinflammation, as shown in pre-clinical studies, possibly through its role in increasing blood brain barrier permeability and attracting monocytes to the brain^35,36^. Thus, acutely raised CCL2 may indicate ongoing neuroinflammation, as a result of peripheral immune cell entry, that is particularly harmful to white matter. IL16 is produced by CD4+ and CD8+ cells, including microglia in the brain, and acts as a chemokine and activating signal for cells expressing the CD4 receptor, such as T-lymphocytes, monocytes and macrophages^37–39^. Peripheral CD4+ T-lymphocyte activation is reported in acute human TBI^21^, while experimental studies have found that CD4+ T-lymphocytes can increase injury severity^40^. Increased IL16 expression is seen in experimental neuroinflammation, as a result of infiltrating immune cells and microglial activation^41^. Our results are thus in keeping with TBI triggering release of DAMPs (damage associated molecular patterns) that cause activated microglial to release IL16, which results in increased immune cell infiltration into the brain and acute neuroinflammation. In keeping with this interpretation, acutely increased CCL2 correlated with lower white matter integrity in the corpus callosum subacutely, a marker of traumatic axonal injury. We have previously shown that axonal damage relates to cognitive and functional outcomes^42–45^, tau deposition^46^ and brain atrophy^47^ after TBI.

There were also TBI-specific changes in other proteins implicated in post-injury neuroinflammation. SFRP1 is a cell-cell signalling molecule, involved in multiple pathways relevant to neuroinflammation, such as *Wnt* signalling. Astrocytic derived SFRP1 after experimental TBI leads to sustained microglial activation, thus promoting a state of chronic neuroinflammation^48^. The clinical significance of our novel finding, that of reduced plasma levels acutely in human TBI, remains to be defined. FCN2 aids clearance of dying cells and is also an initiator of the lectin complement pathway^10,49^, which is activated in brain tissue after experimental TBI^10^. Complement proteins have also been noted to correlate with measures of blood brain barrier permeability in acute severe TBI^50^. The positive correlation with Abeta42 and inverse correlation with pTau231 may indicate that acute activation of these pathways is beneficial after TBI, but this requires further investigation. Sequestosome1 (SQSTM1, also referred to as p62) mediated autophagy^51^ is important for peripheral myeloid cell differentiation^52^ and microglial functions, including degradation of amyloid plaques^53^. Raised SQSTM1 in acute TBI was associated with less axonal injury, as measured on DTI, supporting a role for post-injury autophagy in mitigating axonal injury. Indeed, experimental TBI studies have shown that impaired autophagy by microglial and macrophages exacerbates worse post-injury neuroinflammation, through reduced clearance of DAMPs and proinflammatory signals, such as the NLRP3 inflammasome, and worsens outcomes^9^.

Cell metabolism changes are also important in TBI pathophysiology^54^. Experimental studies have found reduced astrocytic malate dehydrogenase 1 (MDH1) expression in concussive TBI models^55^ and increased thalamic expression in blast TBI models^56^. Acetylation of MDH1 reduces oxidative stress after intracerebral haemorrhage, and reduced MDH1 activity is associated with cell senescence^57,58^. Oxidative stress and abnormal glucose metabolism contributes to secondary injury after the initial TBI event^59,60^. Our study finds that plasma MDH1 levels are specifically increased after TBI, which may reflect post-TBI disruptions in glucose metabolism and response oxidative stress.

Several of the TBI-specific proteins we identified are associated with neurodegenerative disease. For example, pTau231 was elevated after TBI, and is a sensitive marker of early tau pathology in the brain and increases with amyloid beta deposition in pre-clinical Alzheimer’s disease^61^. We, and others, have found evidence of tau pathology after single-hit moderate-severe TBI, beginning as soon as 1 year post-TBI, compared to age-matched controls^46,62,63^. Higher rates of brain atrophy are seen after TBI^64,65^ and experimental studies have also demonstrated that TBI initiates a cascade of self-propagating tau pathology, in a prion-like manner^63^. Therefore, an increase in pTau231, an abnormal phosphorylated tau isoform, in the acute period is intriguing as it may indicate the beginning of pathological processes that leads to later neurodegeneration. In contrast, acute plasma levels of pTau181 and pTau217 were raised compared to CON but did not reach statistically significance compared to the NTT cohort. There have been no prior clinical acute TBI studies of pTau217, and prior studies of pTau181 in TBI have reported conflicting findings about whether pTau181 is acutely elevated in TBI^66,67^. Given the evidence that chronic traumatic encephalopathy (CTE) has distinct tau pathology from other neurodegenerative diseases such as Alzheimer’s^68,69^, the ability to assess multiple isoforms on the same panel may be useful for future *in vivo* differentiation of CTE.

We also found lower levels of plasma Abeta42 in acute TBI, compared with NTT and CON. Prior studies have found both reduced and increased CSF Abeta42 levels^70–72^ in acute TBI, whilst two prior human TBI studies reported raised plasma Abeta42 levels in acute TBI^71,73^. Plasma and CSF amyloid beta 42 (Abeta42) levels are reduced in Alzheimer’s disease, as lower soluble amyloid isoforms decrease as they condense into amyloid plaques^74,75^. Therefore, our finding of reduced plasma Abeta42 in acute TBI, compared to both CON and NTT, fits in with the idea that TBI triggers neurodegenerative processes. Future studies should aim to elucidate whether acute pTau231 and Abeta42 elevation predicts higher rates of brain atrophy and later neurodegeneration after TBI.

Plasma levels of the neurodegenerative proteins presenilin 1 (PSEN1) and 14-3-3γ levels were specifically raised after TBI, compared to non-TBI trauma, which has not been previously reported in clinical TBI studies. Raised plasma PSEN1 may reflect early abnormal amyloid plaque formation, which has previously been demonstrated to occur within 24h after TBI^76^. Amyloid precursor protein (APP) and enzymes required for its cleavage (specifically BACE-1 and PSEN1) and Abeta are co-located in these abnormal diffuse amyloid plaques early after TBI, accumulate in axons in both human pathological and experimental TBI studies, and inhibition of cleavage activity in experimental TBI reduces injury-induced cell loss and behavoiural deficits^77–79^. Axonal disruption and cell death during TBI may promote abnormal amyloid plaque formation by forcing together all the components necessary for abnormal APP cleavage^76^. Raised CSF levels of 14-3-3ζ have been previously reported in TBI^80^, but raised plasma 14-3-3γ (YWHAG) is reported here for the first time. This brain-enriched intracellular signal transduction has been reported to be a component of pathological protein depositions in several neurodegenerative diseases^81^ and is also used as a marker of neuronal injury in the diagnosis of Creutzfeldt-Jakob disease ^82^. Therefore, raised levels of YWHAG in our acute TBI cohort may reflect neuronal damage, with its implications to be investigated in further work.

Exploratory cluster analysis found that heterogeneity of plasma protein expression patterns reflect injury characteristics. This data-driven approach differentiated TBI from non-injured and non-TBI trauma control groups, as well as identified TBI subgroups that had specific pathological characteristics. These novel findings may indicate that different pathophysiological processes mediate the development of white matter versus lesional injury. For example, TBI subgroups with significantly different lesion volumes had markedly different expression patterns of proinflammatory proteins (e.g. the NLRP3 inflammasome-associated protein IL18^83^), neurotrophins (e.g. BDNF) and proteins associated with neuronal injury an1d neurodegenerative disease (e.g. SOD1, TDP-43 proteins and alpha-synuclein proteins). Additionally, acute levels of neurodegenerative proteins (pTau231, total tau, PSEN1) and the neuronal marker UCHL1 correlated with subacute lesion volume, but not white matter integrity. Future work can seek to disentangle whether and how these proteins and associated pathways contribute to lesion development, and whether there is regional variation in the relationship between white and grey matter measures and plasma protein levels, as we have previously shown with GFAP^84^.

Several proteins were assessed with two different assay types, the Alamar NULISA™ assay and/or ELISA and OLINK® PEA assays. There was generally excellent correlation of assay values for overlapping proteins between the Alamar NULISA™ and OLINK® or ELISA-based platforms, and also agreement between panels about the significance of findings. This overall consistency across platforms increases confidence in the novel findings, and further strengthens the case for NFL, GFAP, S100B, total tau and UCHL1 as biomarkers of TBI and its severity. Important exceptions to the generally good correlation between panels are those proteins where >50% of samples were deemed to be below the level of detection by the OLINK® assay. This suggests that protein changes need to be above a certain threshold to be robust enough to be platform agnostic. This is an important consideration for future work, for example, if using these assays to track response to treatment.

Whilst we show several novel and interesting findings, our work is exploratory, requiring repetition plus mechanistic experiments to investigate why and how the protein patterns identified come about, and the clinical implications. We had a relatively small cohort, with moderate-severe injury and only tested acute samples. Future clinical studies should replicate our results in larger cohorts including repetitive and mild injuries, study the evolution of plasma protein changes over time, and investigate relationships with outcomes and patient factors, such as age and sex. It is not possible using the current techniques to differentiate the tissue or cell provenance of proteins, whether their levels represent release due to cell death or unregulated expression, or whether their derangement contributes to or only accompanies TBI pathology. Paired CSF/plasma studies and studies in which proteins are experimentally upregulated or knocked out would help with this. About half of our NTT group were limb fractures, which is a risk factor for delirium, occurring in ∼10-35% of hip fractures^85,86^. Studies of peri-operative delirium have found increased plasma levels of inflammatory cytokines^87^ and tau^88,89^. This means that while the NTT group enabled us to identify protein responses specific to TBI injury, compared with general injury, the NTT group may not be a fully “brain neutral” control group. However, there are clearly important differences between the pathophysiology of TBI and delirium, for example, demonstrated by the fact that raised CSF and plasma GFAP is a robust finding in TBI but not delirium^88–90^. Our results are therefore still likely to reflect elements of pathophysiology specific to TBI, but future work should take into consideration potential brain effects of different non-healthy control groups. Finally, assays that report in relative units, like the Alamar and OLINK® panels in this study, are more suited to the discovery and hypothesis-generating work that we have done than for direct clinical use. The relative units also means that we were not able to assess agreement between assays, only whether there was good correlation.

In conclusion, we identify novel proteins whose plasma levels are specifically deranged in acute TBI, compared with non-TBI injury and non-injured controls. These proteins are involved in neurodegeneration, cell metabolism, autophagy and inflammation. We also show correlations between inflammatory proteins and those associated with neurodegeneration and injury. Further, we find that patterns of protein expression distinguished subgroups with TBI. We highlight how a multiplex proteomic approach can contribute to pathophysiological classification of TBI, which is a crucial step improved prognostication and identifying treatments.

## Supplementary Material

Supplementary material is available at *Brain* online

## Author contributions (CRediT)

Conceptualization: LML, AH, HZ, DJS

Methodology: LML, AH, HZ, NG, KZ, GB, DJS

Validation: EK, AH, HZ, FM, KZ

Data curation: LML, EK, AH, NG, KZ

Formal analysis: LML, ES, KZ, NG, DL

Investigation: LML, EK, AH, NG, KZ, EG, FM, SM, GB, DJS

Resources: LML, AH, HZ, GB, DJS

Writing – original draft: LML

Writing – review and editing: All authors

Supervision: AH, HZ, GB, DJS

Project administration: KZ, NG, GB, DJS

Funding acquisition: GB, DJS

## Acknowledgements

We thank Alamar for providing complimentary testing.

