## Supplementary material for "High-dimensional proteomic analysis for pathophysiological classification of Traumatic Brain Injury": SI

**SUPPLEMENTARY INFORMATION**

Supplementary Table 1: Full list of proteins on the Alamar NULISA™ CNS Diseases panel

| ACHE | CNTN2 | IL12p70 | NGF | SAA1 |
| --- | --- | --- | --- | --- |
| AGRN | CRH | IL13 | NPTX1 | SFRP1 |
| ANXA5 | CRP | IL15 | NPTX2 | SFTPD |
| APOE | CSF2 | IL16 | NPTXR | SLIT2 |
| APOE4 | CST3 | IL17A | NPY | SMOC1 |
| Abeta38 | CX3CL1 | IL18 | NRGN | SNAP25 |
| Abeta40 | CXCL1 | IL1B (IL1β) | Oligo-SNCA | SNCA |
| Abeta42 | CXCL10 | IL2 | PARK7 | SOD1 |
| BACE1 | CXCL8 (IL8) | IL33 | PDGFRB | SQSTM1 |
| BASP1 | ENO2 | IL4 | PDLIM5 | TAFA5 |
| BDNF | FABP3 | IL5 | PGF | TARDBP |
| CALB2 | FCN2 | IL6 | PGK1 | TEK |
| CCL11 | FGF2 | IL6R | POSTN | TIMP3 |
| CCL13 (MCP4) | FLT1 | IL7 | PRDX6 | TNF |
| CCL17 | FOLR1 | IL9 | PSEN1 | TREM1 |
| CCL2 (MCP1) | GDNF | KDR | pTau-181 | TREM2 |
| CCL22 | GFAP | KLK6 | pTau-217 | UCHL1 |
| CCL26 | GOT1 | MAPT (total tau) | pTau-231 | VCAM1 |
| CCL3 | HBA1 | MDH1 | pTDP43-409 | VEGFA |
| CCL4 | ICAM1 | MME | PTN | VEGFD |
| CD40LG | IFNG (IFN γ) | MSLN | REST | VGF |
| CD63 | IGF1 | NCAM1 | RUVBL2 | VSNL1 |
| CHI3L1 | IGFBP7 | NEFH | S100A12 | YWHAG (14-3-3 γ) |
| CHIT1 | IL10 | NEFL (NFL) | S100B | YWHAZ (14-3-3 ζ) |

Names are as referred to by Alamar Biosciences. This panel is adapted from the commercially available platform^90^.

Supplementary Table 2: Differential Expression co-efficients

Proteins from the Alamar NULISA™ CNS Diseases panel showing significantly different plasma levels between the CON, NTT and TBI groups.

| **Protein** | **Role/ Pathway** | **TBI v CON coefficient** | **TBI v NTT coefficient** | **NTT v CON coefficient** | **F** | **Adjusted *p*** |
| --- | --- | --- | --- | --- | --- | --- |
| Abeta42 | Neurodegenerative disease associated | -0.826 | -0.635 | -0.191 | 11.638 | 0.0001 |
| ACHE | Synaptic biology | -0.640 | -0.332 | -0.308 | 17.865 | 0.0000 |
| BACE1 | Neurodegenerative disease associated | -0.620 | -0.358 | -0.262 | 16.525 | 0.0000 |
| BASP1 | Neuronal development, Multiple CNS/PNS functions | 0.599 | 0.240 | 0.358 | 9.320 | 0.0006 |
| CALB2 | Neuronal | 0.657 | 0.432 | 0.225 | 10.431 | 0.0003 |
| CCL13 | Cytokine/chemokine | -0.614 | -0.219 | -0.396 | 5.581 | 0.0102 |
| CCL2 | Cytokine/chemokine | 0.806 | 0.730 | 0.075 | 6.213 | 0.0066 |
| CCL3 | Cytokine/chemokine | 0.947 | 0.419 | 0.528 | 14.409 | 0.0000 |
| CCL4 | Cytokine/chemokine | 0.714 | 0.263 | 0.451 | 5.638 | 0.0100 |
| CHI3L1 | Cytokine/chemokine | 3.155 | 1.351 | 1.804 | 27.266 | 0.0000 |
| CNTN2 | Neuronal development, Cell Adhesion | -0.949 | -0.014 | -0.935 | 6.741 | 0.0045 |
| CRH | Hormone | -1.980 | -0.764 | -1.216 | 24.366 | 0.0000 |
| CRP | Acute phase | 4.021 | 0.084 | 3.937 | 54.922 | 0.0000 |
| CX3CL1 | Cytokine/chemokine | 0.482 | 0.218 | 0.264 | 6.839 | 0.0042 |
| CXCL8/ IL8 | Cytokine/chemokine | 1.249 | 0.542 | 0.708 | 11.431 | 0.0001 |
| ENO2 | Neuronal | 0.734 | 1.464 | -0.730 | 26.163 | 0.0000 |
| FABP3 | Neurodegenerative disease associated | 2.377 | 1.459 | 0.918 | 20.804 | 0.0000 |
| FCN2 | Immune/inflammation | -0.843 | -0.738 | -0.106 | 7.847 | 0.0020 |
| GFAP | Astroglial biology | 6.644 | 6.357 | 0.286 | 418.745 | 0.0000 |
| GOT1 | Cell protein metabolism, Tumour biology | 0.765 | 0.000 | 0.765 | 5.597 | 0.0102 |
| HBA1 | Vascular biology, Neurodegenerative disease associated | 2.954 | -0.523 | 3.477 | 18.763 | 0.0000 |
| IGF1 | Neuronal development, Hormone | -0.216 | -0.119 | -0.097 | 4.004 | 0.0373 |
| IL10 | Cytokine/chemokine | 1.988 | 0.816 | 1.172 | 17.885 | 0.0000 |
| IL15 | Cytokine/chemokine | 0.968 | 0.314 | 0.654 | 30.162 | 0.0000 |
| IL16 | Cytokine/chemokine | 0.940 | 0.631 | 0.309 | 13.162 | 0.0000 |
| IL18 | Cytokine/chemokine | 0.514 | 1.016 | -0.502 | 6.695 | 0.0046 |
| IL1b | Cytokine/chemokine | 0.691 | 1.305 | -0.615 | 21.619 | 0.0000 |
| IL2 | Cytokine/chemokine | -0.638 | -0.424 | -0.214 | 5.768 | 0.0091 |
| IL33 | Cytokine/chemokine | 2.404 | 1.167 | 1.237 | 45.775 | 0.0000 |
| IL4 | Cytokine/chemokine | -1.242 | -0.282 | -0.959 | 11.866 | 0.0001 |
| IL6 | Cytokine/chemokine | 5.841 | 1.722 | 4.119 | 99.073 | 0.0000 |
| KLK6 | Neurodegenerative disease associated | -0.748 | -0.524 | -0.224 | 17.738 | 0.0000 |
| MAPT/ total TAU | Tau biology | 2.247 | 1.732 | 0.515 | 55.399 | 0.0000 |
| MDH1 | Oxidative stress/ cell energy metabolism | 0.654 | 0.878 | -0.224 | 6.318 | 0.0061 |
| NCAM1/ CD56 | Cell adhesion/ neuronal development | -0.532 | -0.266 | -0.266 | 14.774 | 0.0000 |
| NEFH | Neuronal | 3.890 | 1.962 | 1.928 | 13.355 | 0.0000 |
| NEFL | Neuronal | 3.330 | 2.205 | 1.125 | 67.216 | 0.0000 |
| NPTX2 | Synaptic biology | 0.317 | -0.031 | 0.348 | 4.614 | 0.0227 |
| NPY | Multiple CNS/PNS functions | -0.772 | -0.305 | -0.467 | 8.114 | 0.0016 |
| OligoSNCA | Neurodegenerative disease associated | -0.367 | 1.284 | -1.651 | 3.989 | 0.0373 |
| PARK7 | Neurodegenerative disease associated | 0.639 | 1.248 | -0.608 | 9.058 | 0.0008 |
| PDLIM5 | Cell protein metabolism | -1.152 | -0.722 | -0.430 | 4.541 | 0.0239 |
| PGF | Vascular biology | 0.733 | -0.029 | 0.763 | 22.386 | 0.0000 |
| PGK1 | Oxidative stress/ cell energy metabolism | 0.809 | 1.859 | -1.050 | 7.645 | 0.0023 |
| POSTN | Oxidative stress/ cell energy metabolism, Tissue remodelling | -0.773 | -0.334 | -0.439 | 13.448 | 0.0000 |
| PRDX6 | Oxidative stress/ cell energy metabolism | 0.412 | 0.757 | -0.345 | 5.358 | 0.0122 |
| PSEN1 | Neurodegenerative disease associated | 1.203 | 0.868 | 0.335 | 14.065 | 0.0000 |
| pTau181 | Tau biology | 1.030 | 0.530 | 0.500 | 12.516 | 0.0001 |
| pTau217 | Tau biology | 0.749 | 0.484 | 0.265 | 13.670 | 0.0000 |
| pTau231 | Tau biology | 1.623 | 1.064 | 0.558 | 24.974 | 0.0000 |
| pTDP43.409 | Neurodegenerative disease associated | 0.474 | 1.255 | -0.781 | 6.861 | 0.0042 |
| REST | Transcription factor, Neurodegenerative disease associated | 2.572 | 1.105 | 1.467 | 43.470 | 0.0000 |
| RUVBL2 | Neuronal development, Gene expression | 0.489 | 1.017 | -0.528 | 6.666 | 0.0046 |
| S100A12 | Immune/Inflammation | 1.513 | -0.893 | 2.406 | 20.890 | 0.0000 |
| S100B | Astroglial biology | 2.057 | 1.593 | 0.463 | 56.228 | 0.0000 |
| SAA1 | Acute phase | 5.464 | 0.417 | 5.047 | 75.694 | 0.0000 |
| SFRP1 | Cell-cell signalling, Choroid Plexus | -0.822 | -0.950 | 0.128 | 7.244 | 0.0032 |
| SLIT2 | Neuronal development, Tumour biology | -0.307 | -0.094 | -0.213 | 4.479 | 0.0249 |
| SMOC1 | Brain tumour, Neurodegenerative disease associated | 0.440 | -0.267 | 0.707 | 5.826 | 0.0088 |
| SNAP25 | Synaptic biology | 0.402 | 0.453 | -0.051 | 13.698 | 0.0000 |
| SNCA | Neurodegenerative disease associated | -0.294 | 0.557 | -0.851 | 4.039 | 0.0367 |
| SOD1 | Neurodegenerative disease associated, Oxidative stress/ cell energy metabolism | 0.018 | 0.728 | -0.710 | 4.672 | 0.0219 |
| SQSTM1 | Autophagy | 1.120 | 0.764 | 0.356 | 19.513 | 0.0000 |
| TAFA5 | Cytokine/chemokine | -1.435 | -0.785 | -0.649 | 23.553 | 0.0000 |
| TARDBP/ TDP43 | Neurodegenerative disease associated | 0.460 | 1.136 | -0.675 | 5.875 | 0.0086 |
| TREM1 | Immune/Inflammation | 0.647 | 0.305 | 0.342 | 12.748 | 0.0000 |
| UCHL1 | Neuronal | 0.630 | 0.583 | 0.047 | 35.820 | 0.0000 |
| VEGFA | Vascular biology | 0.307 | 0.191 | 0.116 | 5.285 | 0.0129 |
| VEGFD | Vascular biology, Tumour biology | -0.176 | -0.355 | 0.179 | 6.841 | 0.0042 |
| VSNL1 | Neuronal | 1.880 | 1.517 | 0.363 | 26.587 | 0.0000 |
| YWHAG/ 14-3-3 | Neurodegenerative disease associated | 0.239 | 0.251 | -0.012 | 6.024 | 0.0076 |

CON = non-injured controls, NTT = non-TBI injury controls, TBI = traumatic brain injury. Negative coefficient denotes where plasma levels in first named group is less than in the second named group (i.e. in the first column, a negative coefficient denotes plasma levels in TBI group are lower than in CON group).

Supplementary Figure 1:

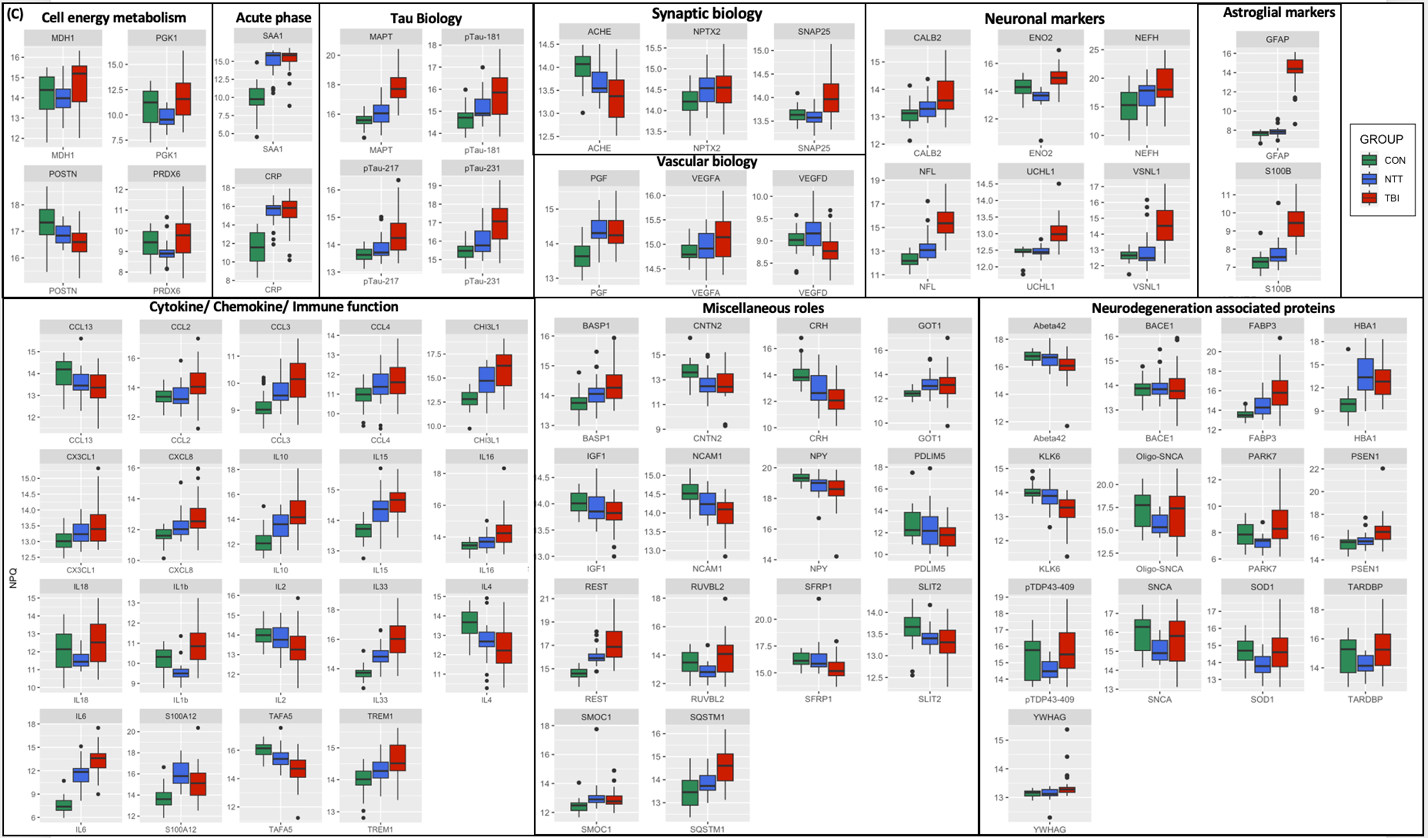

*SI Figure 1: Boxplots illustrating plasma protein levels in CON, NTT and TBI groups for proteins identified by DE analysis to have significant group difference, irrespective of size of comparison coefficient. Y-axis units are NPQ.*

Supplementary Figure 2:

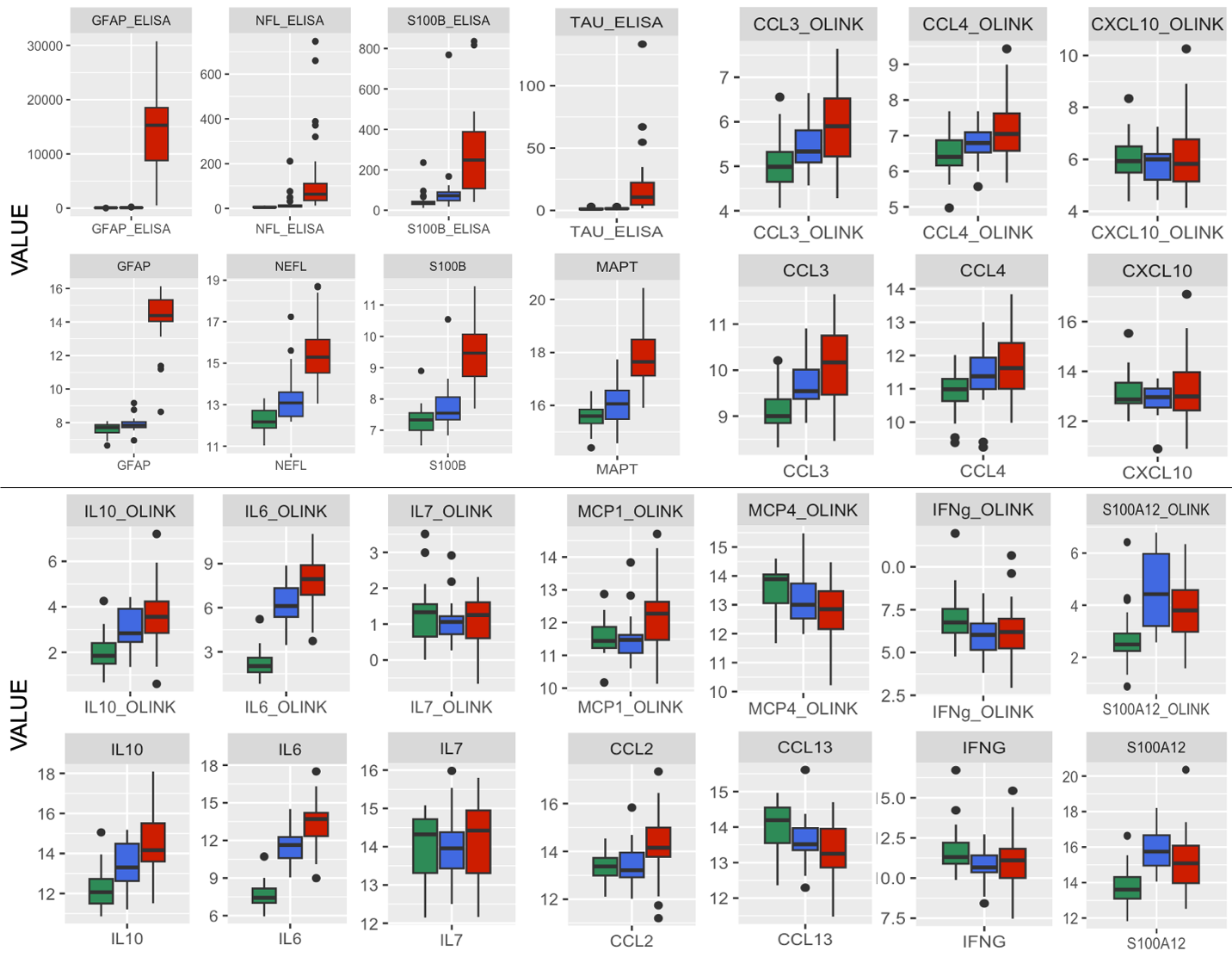

*SI Figure 2: Plots denoting plasma protein levels for each group (TBI=traumatic brain injury [red], CON=non-injured healthy control [green], NTT=non-TBI trauma [blue]). Top row within each panel are from Simoa® (NFL, GFAP, Tau/MAPT) or Millipore (S100B) ELISA-based or OLINK® Target 96 Inflammation assays. Bottom row within each panel is from the Alamar NULISA™ CNS Diseases assay.*

Supplementary Table 3: ANOVA test statistics for overlapping proteins

Statistical tests (ANOVA) of effect of group, and post-hoc (Tukey HSD test) tests, for proteins which overlapped two assay platforms, and where the correlation coefficient between the two platforms >80%.

| **Protein** | **Alamar ANOVA**  (F, *p*) | **Alamar post-hoc NTTvCON**  (t statistic, *p*) | **Alamar post-hoc TBIvCON**  (t statistic, *p*) | **Alamar post-hoc TBIvNTT**  (t statistic, *p*) | **OLINK/ ELISA ANOVA**  (F, *p*) | **OLINK/ELISA post-hoc NTTvCON**  (t statistic, *p*) | **OLINK/ELISA post-hoc TBIvCON**  (t statistic *p*) | **OLINK/ELISA post-hoc TBIvNTT**  (t statistic, *p*) |
| --- | --- | --- | --- | --- | --- | --- | --- | --- |
| GFAP | 390.7,  *<2e^-16^* | 0.2988494, *0.6314965* | 6.7147284, *0.0000000* | 6.4158790, *0.0000000* | 59.28,  *3.06e^-16^* | 42.83789, *0.9997224* | 15048.91667, *0.0000000* | 15006.07878, *0.0000000* |
| S100B | 50.49,  *1.12e^-14^* | 0.5017895, *0.1389913* | 2.0753420, *0.0000000* | 1.5735525, *0.0000000* | 15.97,  *1.62e^-06^* | 62.85347, *0.4158558* | 223.85212, *0.0000023* | 160.99865, *0.0020133* |
| NFL/ NEFL | 56.84,  *8.05e^-16^* | 1.194252, *0.0064301* | 3.299303, *0.0000000* | 2.105051, *0.0000002* | 8.976, *0.000317* | 22.15665, *0.8184366* | 120.76415, *0.0005670* | 98.60750, *0.0127923* |
| Total TAU/ MAPT | 53.61,  *3.01e^-15^* | 0.5387181,  *0.1345295* | 2.2731316,  *0.0000000* | 1.7344135,  *0.0000000* | 9.606,  *0.000191* | 0.2430742,  *0.9988068* | 16.6585538,  *0.0008549* | 16.4154796,  *0.0029647* |
| CCL3 | *12.95,*  *1.45e^-05^* | 0.5239004, *0.0558309* | 0.9544832, *0.0000078* | 0.4305828, *0.0986301* | 9.529, *0.000203* | 0.4319678, *0.1507354* | 0.8387072, *0.0001235* | 0.4067394, *0.1391207* |
| CCL4 | 5.337, *0.00678* | 0.4029678, *0.2905964* | 0.7303875, *0.0046663* | 0.3274197, *0.3810165* | 7.03, *0.00158* | 0.3795537, *0.2045105* | 0.6944522, *0.0010133* | 0.3148984, *0.2745772* |
| CXCL10 | 0.701,  *0.499* | -0.2876273, *0.6334473* | 0.0482560, *0.9818416* | 0.3358833, *0.4820932* | 0.517,  *0.599* | -0.32200528, *0.6060549* | -0.05470318, *0.9795881* | 0.26730210, *0.6657029* |
| MCP1/CCL2* | 6.116, *0.00345* | 0.07878943, *0.9680564* | 0.85268556, *0.0070270* | 0.77389613, *0.0307995* | 4.257, *0.0177* | 0.01275153, *0.9987902* | 0.57958276, *0.0352150* | 0.56683123, *0.0677883* |
| MCP4/CCL13 | 5.153, *0.00797* | -0.3552136, *0.2719566* | -0.6165734, *0.0054838* | -0.2613598, *0.4332453* | 6.026, *0.00372* | (-0.4132767, *0.2812844* | -0.7842168, *0.0025246* | -0.3709401, *0.2999099* |
| IFNg* | 3.469, *0.0362* | -1.1077335, *0.0495615* | -0.8422095, *0.0837780* | 0.2655240, *0.8081663* | 4.38,  *0.0158* | -1.1709607, *0.0308296* | -0.9728204, *0.0332727* | 0.1981403, *0.8833220* |
| IL6 | 92.5  *<2e^-16^* | 3.942867, *0.0000000* | 5.757023, *0.0000000* | 1.814156, *0.0006028* | 92.4,  *<2e^-16^* | 4.05335, *0.0000000* | 5.59463, *0.0000000* | 1.54128, *0.0031231* |
| IL7 | 0.371,  *0.692* | -0.02474654, *0.9961694* | 0.17315439, *0.7673885* | 0.19790092, *0.7500838* | 0.273,  *0.762* | -0.15781875, *0.7581300* | -0.09995857, *0.8542463* | 0.05786018, *0.9569621* |
| IL10 | 15.54,  *2.2e^-06^* | 1.1272697, *0.0258349* | 1.9878442, *0.0000011* | 0.8605745, *0.0777274* | 12.99,  *1.4e^-05^* | 1.0123571, *0.0187631* | 1.5611849, *0.0000074* | 0.5488278, *0.2381324* |
| S100A12 | 14.85,  *3.59e^-06^* | 2.1782513, *0.0000056* | 1.4634882, *0.0002707* | -0.7147631, *0.1638905* | 13.43,  *1.02e^-05^* | 1.9449921, *0.0000079* | 1.1205100, *0.0023812* | -0.8244821, *0.0569925* |

* denotes proteins where the post-hoc tests would lead to different conclusions about the relative effect of group on protein levels. CON=non-injured healthy controls, NTT=non-TBI trauma injury, TBI=traumatic brain injury. GFAP, NFL and S100B were assayed on ELISA-based platforms in addition to the Alamar platform, whilst the cytokines were assayed on the OLINK® platform in addition to the Alamar platform.
